# Biophysical analysis of *Plasmodium falciparum* Hsp70-Hsp90 organizing protein (PfHop) reveals a monomer that is characterised by folded segments connected by flexible linkers

**DOI:** 10.1101/866137

**Authors:** Stanley Makumire, Tawanda Zininga, Juha Vahokoski, Inari Kursula, Addmore Shonhai

**Affiliations:** Department of Biochemistry, School of Mathematical & Natural Sciences, University of Venda, Thohoyandou, South Africa; Department of Biomedicine, University of Bergen, Bergen, Norway; Biocenter Oulu & Faculty of Biochemistry and Molecular Medicine, University of Oulu, Oulu, Finland

## Abstract

*Plasmodium falciparum* causes the most lethal form of malaria. The cooperation of heat shock protein (Hsp) 70 and 90 is important for folding of a select number of cellular proteins that are crucial for cyto-protection and development of the parasites. Hsp70 and Hsp90 are brought into a functional complex that allows substrate exchange by stress inducible protein 1 (STI1), also known as Hsp70-Hsp90 organizing protein (Hop). *P. falciparum* Hop (PfHop) co-localises and occurs in complex with the parasite cytosolic chaperones, PfHsp70-1 and PfHsp90. Here, we characterised the structure of recombinant PfHop using synchrotron radiation circular dichroism (SRCD) and small-angle X-ray scattering. Structurally, PfHop is a monomeric, elongated but folded protein, in agreement with its predicted TPR domain structure. Using SRCD, we established that PfHop is unstable at temperatures higher than 40 °C. This suggests that PfHop is less stable at elevated temperatures compared to its functional partner, PfHsp70-1, that is reportedly stable at temperatures as high as 80 °C. These findings contribute towards our understanding of the role of the Hop-mediated functional partnership between Hsp70 and Hsp90.

## Introduction

Heat shock proteins (Hsp) serve primarily as protein folding facilitators. They also participate in several other processes, such as protein transport, assembly/disassembly of protein complexes, protein degradation, amongst others [1]. Their role in the survival and pathogenicity of malaria parasites is increasingly becoming apparent [2; 3; 4]. *Plasmodium falciparum* is the agent for the most lethal form of malaria. It has been reported that the cytosolic *P. falciparum* heat shock protein 70-1 (PfHsp70-1) is cyto-protective due to its ability to suppress protein mis-folding and aggregation under stressful conditions [5; 6]. In addition, another cytosolic molecular chaperone, *P. falciparum* Hsp90 (PfHsp90) is essential [7]. The cooperation of Hsp70 and Hsp90 is known to facilitate folding and function of proteins implicated in cell development, such as steroid hormone receptors and kinases [8; 9].

Stress inducible protein 1 (STI1) was first described in mouse [10], and now also known as Hsp70-Hsp90 organizing protein (Hop), acts as a module that allows Hsp70 and Hsp90 to interact stably, thereby facilitating substrate transfer from Hsp70 to Hsp90. Hop is conserved and stress inducible protein that possesses three tetratricopeptide repeats (TPR): TPR1, TPR2A and TPR2B [10]. Both Hsp70 and Hsp90 interact with Hop via the C-terminal EEVD motif, present in the two molecular chaperones [11; 12]. Hop interacts with Hsp70 and Hsp90 via its TPR1 and TPR2A domains, respectively [11]. While for a long time the role of the TPR2B domain of Hop has remained largely elusive, it is now thought that Hsp70 first binds to the TPR1 domain of Hop before switching to the TPR2B domain to facilitate substrate transfer to Hsp90 [11; 13].

In light of the importance of both PfHsp70-1 and PfHsp90 in the survival of the malaria parasite, there has been growing interest in identifying inhibitors targeting the function of these two molecular chaperones. Compounds that inhibit PfHsp70-1 [14; 15; 16] and PfHsp90 [17, 9] have been identified, and some of them exhibit antiplasmodial activity. Some compounds that target PfHsp90 function reverse parasite resistance to traditional antimalarial drugs, such as chloroquine (reviewed in [18]). We previously described *Plasmodium falciparum* Hop (PfHop), which co-localises and associates with both PfHsp70-1 and PfHsp90 [19; 12]. While Hop in other organisms, such as yeast and human, has been rather extensively characterised, the structure and function of PfHop remain to be elucidated.

Here, we show that PfHop is a monomeric, elongated but folded protein, which loses most of its secondary structure at temperatures above 40°C. We discuss the implications of our findings with respect to the role of PfHop in coordinating the Hsp70-Hsp90 pathway in *P. falciparum*.

## 2.0 Methods and Materials

### 2.1 Materials

Reagents used in this study, unless otherwise stated, were purchased from Merck Chemicals (Darmstadt, Germany), Thermo Scientific (Illinois, USA), Zymo Research (USA), Melford (Suffolk, UK), and Sigma-Aldrich (USA). Nickel NTA resin was purchased from Thermo Scientific (USA). ECL was purchased from (ThermoFisher Scientific, USA). The expression and purification of his-tagged recombinant forms of PfHop was confirmed by Western blotting using anti-His antibodies (Thermo Scientific, USA). Furthermore, rabbit raised anti-PfHop antibodies (Eurogentec, Belgium; 19) were also used to confirm the presence of recombinant PfHop protein.

### 2.2 Expression and purification of recombinant PfHop

Recombinant PfHop (PF3D7_1434300) was overexpressed in *Escherichia coli* XL1 Blue cells and purified by nickel affinity chromatography as previously described [19; 12]. The Ni-affinity purified proteins were extensively dialysed in SnakeSkin dialysis tubing 10 000 MWCO (ThermoFisher Scientific, USA) against buffer A [20 mM Tris-HCl, pH 7.5, 10 mM NaCl, 5% (v/v) glycerol, 0.2 mM Tris-carboxyethyl phosphine (TCEP)]. The protein from Ni-NTA chromatography was used to perform conventional CD spectroscopy and tryptophan fluorescence assays. SRCD and SAXS analyses were performed using protein that was further purified using ion exchange and size exclusion chromatography as follows. The dialysed protein was further purified using anion exchange chromatography using a Tricorn MonoQ 4.6/100 PE column (G.E Healthcare LS, USA). PfHop was eluted by applying buffer B (20 mM Tris-HCl, pH 7.5, 10 mM NaCl, 0.2 mM TCEP) to the column using a linear (0.1-1.0 M) NaCl gradient. As the final purification step and to evaluate the oligomeric state of PfHop, size exclusion chromatography was used. Following anion exchange, fractions containing pure PfHop were pooled together and loaded onto a HiLoad 16/600 Superdex™ 200 pg column equilibrated with buffer C (10 mM Tris-HCl, pH 8, 300 mM NaCl containing 5% glycerol, 0.2 mM TCEP). Eluted fractions were analysed using SDS-PAGE to determine the purity and homogeneity of the PfHop protein. Authenticity of the purified protein was confirmed by sequencing using MALDI-TOF mass spectrometry at the Biocenter Oulu Proteomics Core Facility, Oulu University, Finland. The protein concentration was determined by measuring the UV absorbance at 280 nm using a Nanodrop ND100 (ThermoFisher Scientific, USA).

The molecular weight of PfHop was determined using multi-angle static light scattering (MALS) coupled to size exclusion chromatography using a Superdex S200 Increase 10/300 GL column (GE Healthcare). The column, equilibrated with buffer C, was coupled to a mini DAWN TREOS MALS detector (Wyatt Technology, Germany) and an ERC RefraMAx520 differential refractometer (ERC, Germany). 100 µg of PfHop in buffer C was injected into the column using a flow rate of 0.5 ml/min. BSA and ovalbumin were used as molecular-weight controls. The molecular weight of PfHop was determined based on the measured light scattering at three different angles and the refractive index using the ASTRA software version 6.1.5.22 (Wyatt Technology, Germany).

### 2.4 Investigation of the secondary structure of PfHop

The secondary structure of PfHop was investigated using synchrotron radiation (SR) and conventional circular dichroism (CD) spectroscopy. The spectral measurements were conducted at the UV-CD12 beam line (Anka, Karlsruhe) under temperature-controlled conditions. PfHop at a concentration of 0.5 mg/ml dialysed in buffer D (10 mM K_3_PO_4_, pH 7.0, 150 NaF) was analysed using a 98.56 µm path length round cell cuvette (Suprasil, Hellma Analytics, Germany) at a constant temperature of 10°C. A total of 3 full spectral scans were recorded from 280 to 175 nm and averaged. CD spectroscopy experiments were done using a Jasco J-1500 CD spectrometer (JASCO Ltd, UK) with a temperature-controlled Peltier. Recombinant proteins at a final concentration of 2 μM were analysed using a 2-mm path-length quartz cuvette (Hellma). Spectral scans were recorded from 250 to 180 nm and averaged for least 3 scans. The SRCD data were processed and deconvoluted using the Dichroweb server (20) and the CONTINLL algorithm with the SP175 reference set (21). To further predict the secondary structure content of PfHop, the BeStSel server (http://bestsel.elte.hu/; [22; 23]) and the Phyre2 server (http://sbg.bio.ic.ac.uk/phyre2/; [24]) were also used. In order to investigate the heat stability of PfHop, the protein was subjected to increasing temperature (10 to 90°C using 5°C intervals) and full spectrums recorded. Melting curves were plotted by monitoring the CD signal at 192, 193, 210 and 220 nm. The measurements were expressed as folded protein fraction at the respective temperature using previously described protocols [25; 26]. The tertiary structure of the protein was probed in the presence of varying concentrations of denaturants, urea (0 - 8 M) and guanidine hydrochloride (0 - 6 M).

Fluorescence spectra were recorded with initial excitation at 295 nm and emission being measured between 300 nm and 400 nm using JASCO FP-6300 spectrofluorometer (JASCO, Tokyo, Japan).

### 2.5 Small-angle X-ray scattering analysis for PfHop shape determination

Synchrotron small-angle X-ray scattering (SAXS) data were collected on the EMBL Hamburg Outstation beam line P12 at PETRA III/DESY (Hamburg). PfHop (2.2 mg/ml) and buffer samples were exposed to X-rays with a wavelength of 1.240 Å for 0.045 s. Pre-processed data were further analysed with the ATSAS software package [27]. The distance distribution calculation and *ab initio* modeling were performed using GNOM [28] and the GASBOR package [29], respectively. Human Hop Tpr1 (1ELW; 0) and bakers’s yeast Tpr2AB domains (3QU3; 11) were manually fitted in the *ab initio* envelope using PyMOL 2.3.2 (Schrödinger, USA). An R_g_ value of 5.3 nm was determined visually from the linear part of the low scattering angles (0.0087 – 0.0715 nm^-1^) using PRIMUS (1). The D_max_ for PfHop was estimated as 24 nm, also using PRIMUS.

## 3 Results

### 3.1 Oligomeric state of recombinant PfHop

Recombinant PfHop was purified using nickel affinity chromatography as previously described [19]. The protein was further purified using ion exchange and subsequently subjected to size exclusion chromatography (Supplementary Figure S1).

Hop has been reported to exist as either monomer [32] or dimer [33], or is largely monomeric forming weak dimers [34]. Based on size exclusion chromatography, PfHop eluted as an elongated monomer under reducing conditions. A small fraction of dimer, likely due to partial oxidation, could be seen in some batches (Figure S1 C, lanes 1-5). The monomeric state was confirmed using multi-angle light scattering coupled to size exclusion chromatography, which gave a molecular weight of 73 kDa for the main peak (Figure 1). This is 8% larger than the calculated theoretical molecular weight 67.6 kDa. As a control, we determined molecular weights for bovine serum albumin (73 kDa) and ovalbumin (45 kDa), which were 10% and 6%, respectively, larger than their calculated theoretical molecular weights.

**Figure 1.**
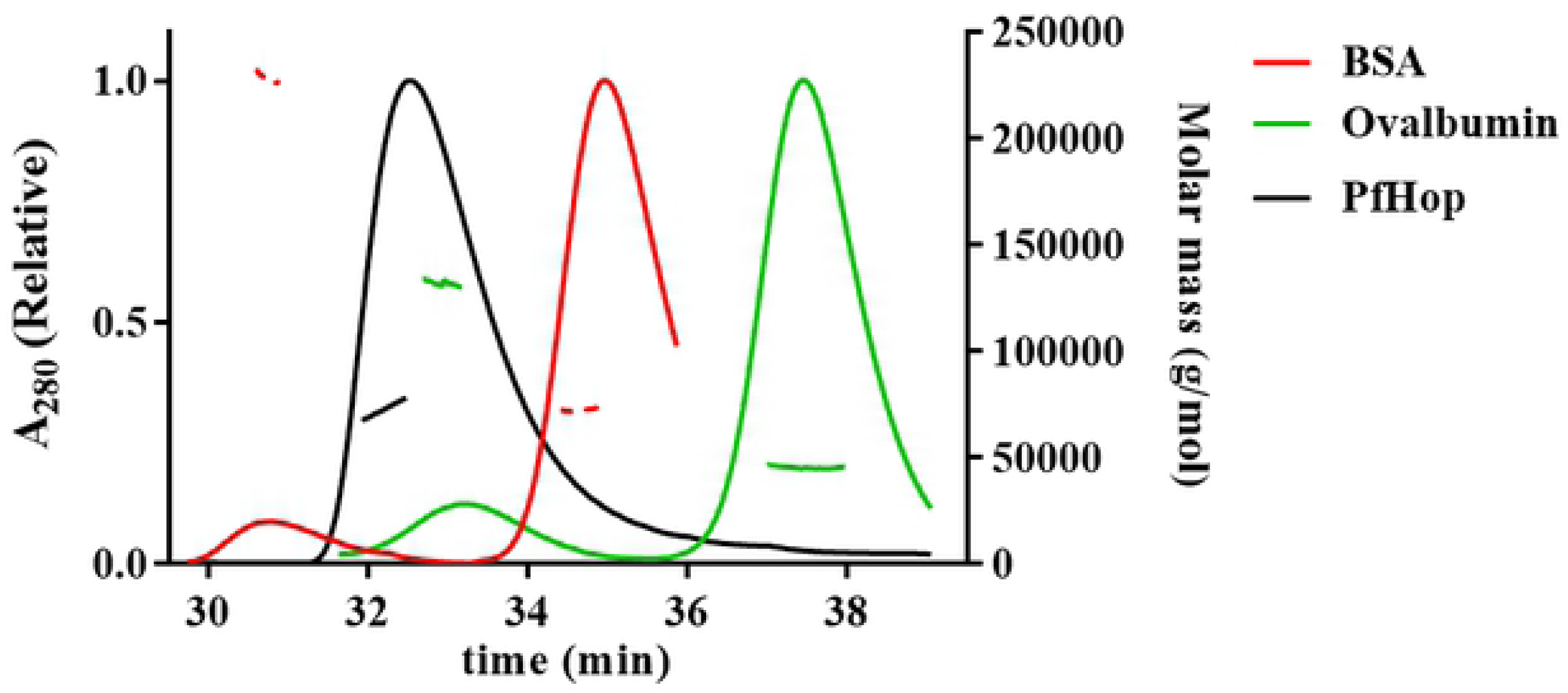
Determination of the oligomeric status of PfHop. Size exclusion chromatography of PfHop displays a single peak. The molecular weight of the peak calculated using static light scattering, shown as a black line, represents a PfHop monomer.

### 3.3 Analysis of the secondary structure of PfHop

To confirm the folding state and secondary structure composition of recombinant PfHop, synchrotron radiation circular dichroism (SRCD) spectroscopy was conducted. SRCD spectra were recorded between 175 and 280 nm at 10°C. The PfHop spectra exhibited 2 negative minima around 222 and 208 nm and a positive peak at 194 nm (Figure 2A), characteristic of a predominantly α-helical protein [35]. Deconvolution of the spectra with Dichroweb indicated a predominantly α-helical structure comprising 77 % α-helices (Supplementary Table 1). This was supported by predictions from BeStSel and Phyre2 (Supplementary Table 1). The observed predominantly α-helical structure is consistent with the predicted three-dimensional model of PfHop which showed that all its three TPR motifs are α-helical in nature [19]. Furthermore, based on the previously generated three-dimensional model of PfHop, residues of PfHop that are implicated in making direct contact with PfHsp70-1/PfHsp90 are positioned within the grooves of the α-helical TPR domains [19]

**Figure 2.**
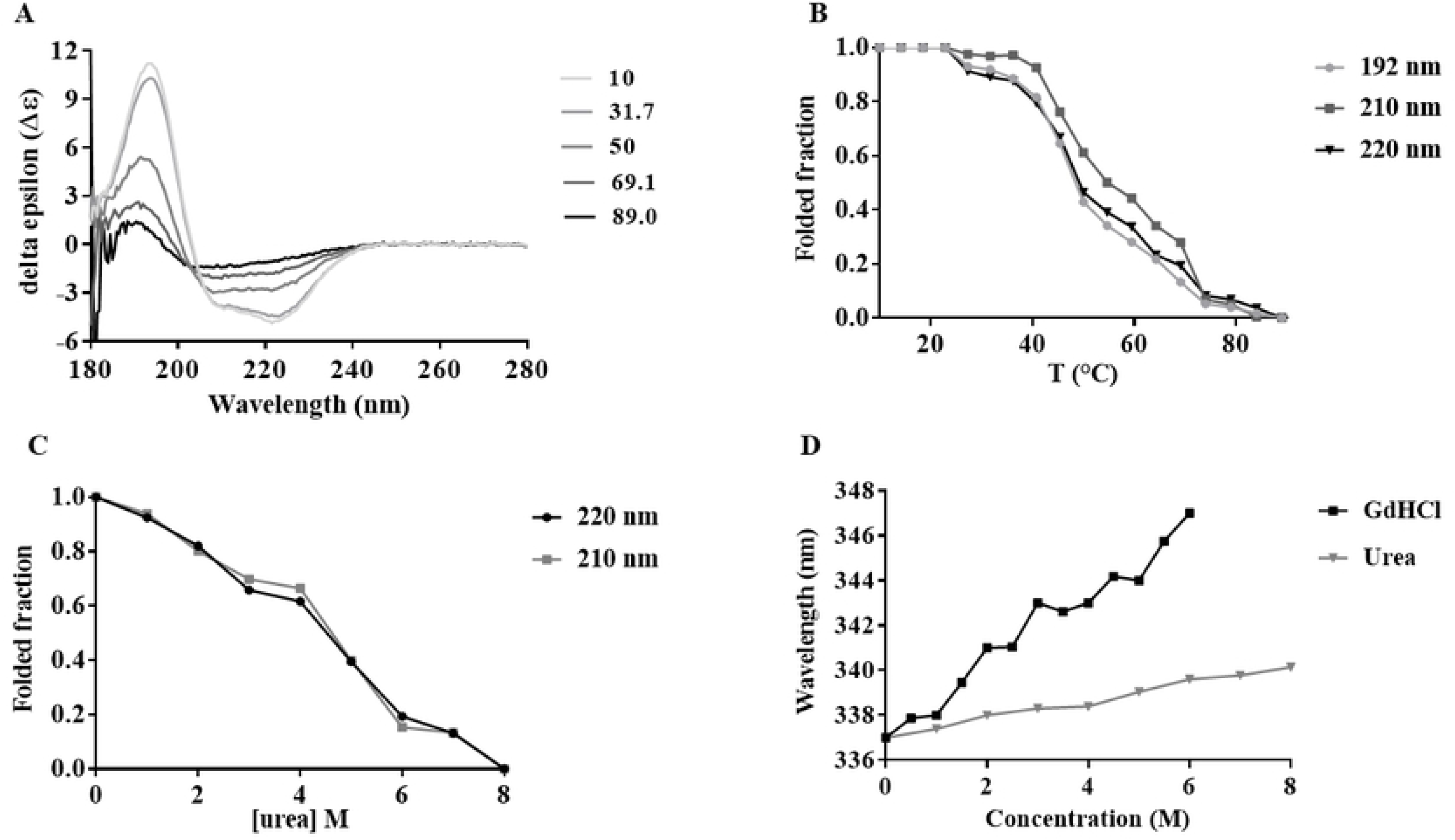
Secondary structure analysis of PfHop. (**A**) SRCD spectrum of full-length PfHop. SRCD spectral scans upon heating from 10 to 90 °C, resulting in unfolding of PfHop. (**B**) The folded fraction of PfHop as a function of temperature. (**C)** Urea-induced unfolding of PfHop. (**D**) The fluorescence emission spectra monitored at 300-450 nm after an initial excitation at 295 nm. The recombinant PfHop protein tryptophan fluorescence emission spectra were recorded under various GdHCl and urea concentrations. Assessment of the red shift of PfHop exposed to different GdHCl and urea concentrations on the emission spectra were plotted.

PfHop mediates the interaction between PfHsp70-1 and PfHsp90 [19; 12], and its expression is heat-induced [19]. The roles of these two chaperones become important when the parasite is under heat stress, such as during clinical malaria fever episodes [36]. It is therefore important that heat shock proteins of parasite origin exhibit resilience to heat stress conditions, and PfHsp70-1 is stable at high temperatures and is most active at 48-50 °C and retains its ATPase activity at up to 80 °C [6; 25]. However, it remains to be established whether PfHop exhibits the same resilience to heat stress.

To probe this, we investigated the heat stability of recombinant PfHop in vitro. As a control, the denaturation of PfHop exposed to urea (0 - 8 M) was also monitored using CD (Figure 2C). Next, we monitored the folded fraction of PfHop in response to exposure to increased temperature conditions using SRCD (10 to 90 °C) (Figure 2A). PfHop appeared stable at temperatures lower than 40°C. However, at higher temperatures the protein lost its fold, and only 50% of the protein retained its folded state at 45 °C (Figure 2B). Notably, the spectra suggest that the protein simultaneously loses both its α-helical and β-compositions in response to heat stress. Based on these findings, PfHop is less stable to heat stress than PfHsp70-1, whose ATPase activity was found to be optimal at 50 °C [25] and exhibits chaperone activity (suppressing heat induced aggregation of protein) at 48 °C [6].

Furthermore, tryptophan fluorescence spectroscopy was conducted to monitor the tertiary structural organization of PfHop in the presence of varying amounts of urea and guanidine hydrochloride (GdHCl). A red shift was observed with maximum peaks at 350 nm (associated with 6 M GdHCl) and 343 nm (associated with 8 M urea) (Figure 2D). PfHop was more sensitive to GdHCl, which is a stronger denaturant. This is in agreement to a previous observation for PfHsp70-1 protein [25].

### 3.4 Low-resolution structure of PfHop in solution

In order to gain further insight into the structure of PfHop, we determined its low-resolution structure in solution using SAXS (Figure 3). The X-ray scattering curve (Figure 3A), the Kratky plot (Figure 3B), and the distance distribution function (Figure 3C) together indicate that PfHop is an elongated protein with a maximum dimension of approximately 24 nm (Table 1). PfHop consists mostly of folded parts connected by flexible linkers. Thus, as expected, the TPR domains likely are arranged like pearls on a string. The distance distribution function (Figure 3C) indicates at least two stable domains with maxima at ∼3 and ∼5 nm within the D_max_ of 24 nm. An *ab initio* dummy residue model calculated with GASBOR (Figure 3D) is consistent with the above and shows an excellent fit to the experimental data (Figure 3) with a χ^2^ value of 1.06. The model has an elongated shape with some more compact regions, to which crystal structures of human and yeast Hop TPR1 (1ELW, 30) and TPR2AB (3UQ3, 11) domains, respectively, fit visually well (Figure 3D).

**Table 1.**
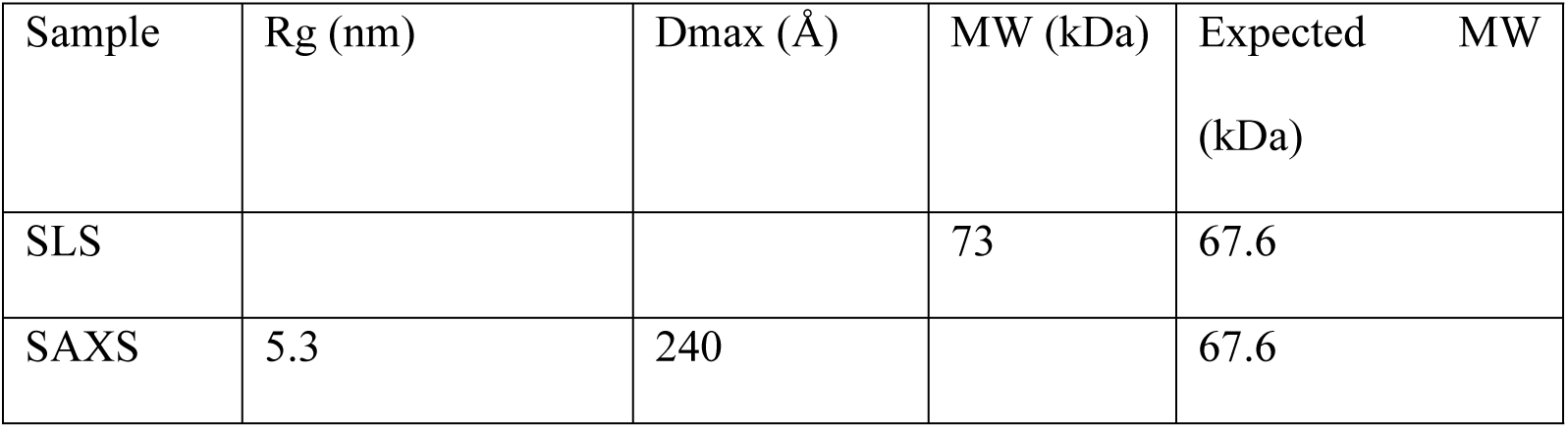
SAXS parameters for PfHop.

**Figure 3.**
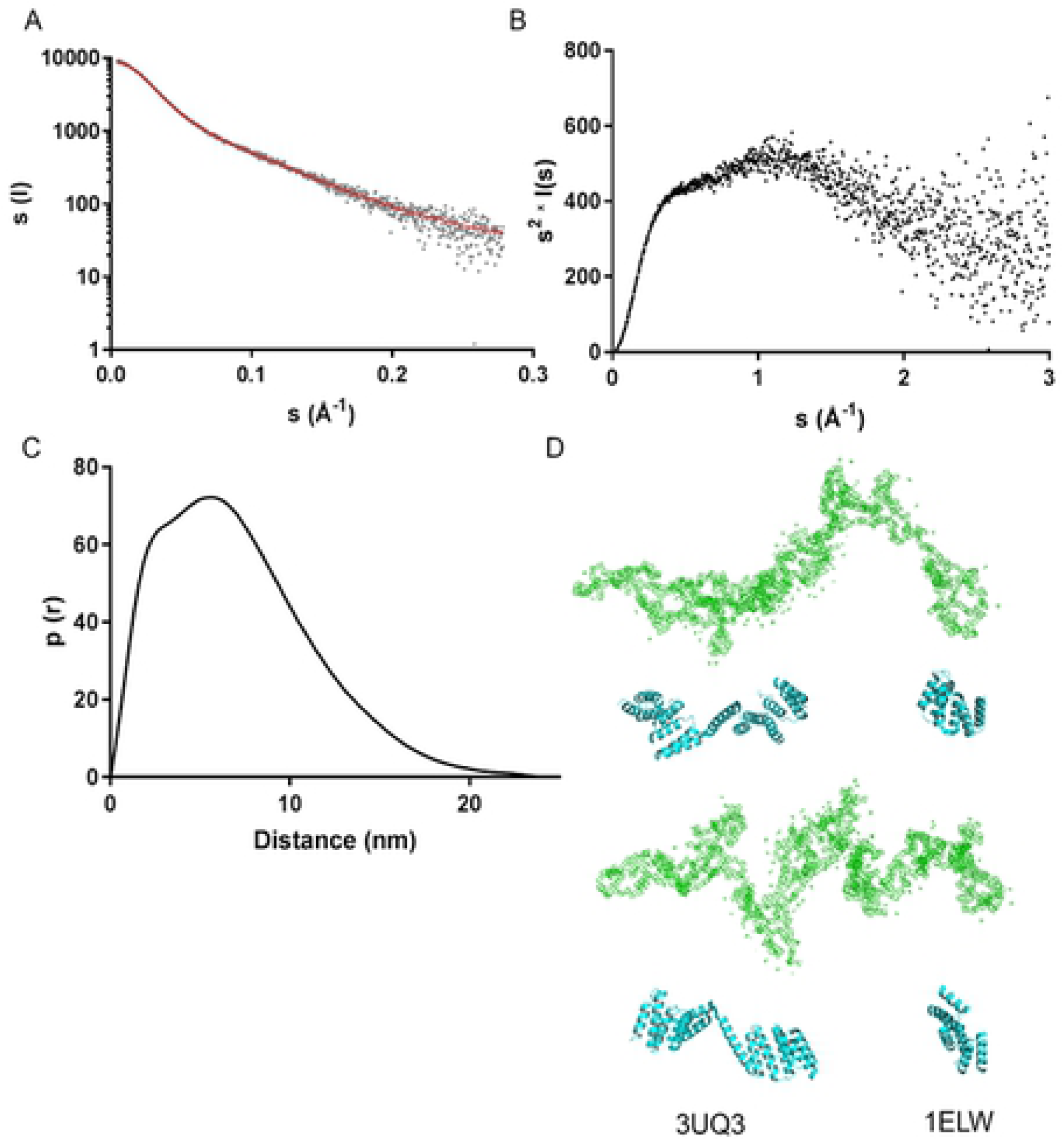
SAXS analysis of PfHop. (A) Fit of a calculated SAXS curve based on an *ab initio* model (red line) on the experimental SAXS curve (grey dots) measured for PfHop. (B) Kratky plot derived from the scattering data. (C) Distance distribution function. (D) An *Ab initio* model of PfHop (green), determined using GASBOR compared with crystal structures of human Hop TPR2AB (3UQ3) and baker’s yeast TPR1 domains (1ELW) (cyan). The lower panel is related to the upper one, by a 90° clockwise rotation along the plane of view.

## 4. Discussion

PfHop is thought to facilitate the functional cooperation between PfHsp70-1 and PfHsp90, both prominent cytosolic molecular chaperones of *P. falciparum* [19; 12]. The Hsp70-Hop-Hsp90 pathway plays an important role in cellular development, as it facilitates folding and maturation of proteins, such as kinases and steroid hormone receptors [37]. The inhibition of both PfHsp90 and PfHsp70-1 leads to parasite death [7; 14; 16], making these molecular chaperones potential antimalarial drug targets. In the current study, we demonstrated that PfHop PfHop is unstable at temperatures above 40 °C (Figure 2). We previously observed that PfHop, PfHsp70-1, and PfHsp90 occur in a complex, and that PfHop directly associates with the TPR domains of PfHsp70-1 and PfHsp90 [19; 12]. This suggests that PfHop modulates functional cooperation between PfHsp70-1 and PfHsp90. The development of clinical malaria is associated with body temperature rising to 41.6 °C. At such high temperatures, the role of heat shock proteins in maintaining proteostasis becomes unquestionably important, as is evidenced by the elevated expression of key molecular chaperones, such as PfHsp70-1 and PfHsp90 [7; 38]. It is intriguing to imagine how PfHsp70-1 and PfHsp90 cooperate in the presence of limiting levels of PfHop, as occurs during sustained heat stress conditions induced by malaria fever. It is possible that under extended stress conditions, the role of Hop becomes less vital and that perhaps Hsp70 and Hsp90 may directly interact. Indeed, a study showed that despite lack of Hop in *E. coli*, Hsp70 and Hsp90 from *E. coli* are capable of direct interaction [39]. In a previous study, we observed complexes of PfHsp70-1 and PfHsp90 in which PfHop was present based on size exclusion chromatography of parasite lysates [19]. However, we also observed eluates representing a complex of PfHsp70-1 and PfHsp90, in which PfHop was absent [19]. In addition, yeast Hsp70 and Hsp90 were recently shown to directly bind in the absence of Hop [40]. A non-canonical Hop homologue from *Caenorhabditis elegans* has been shown to lack a TPR1 domain, thus was shown to be more biased towards binding to Hsp90 than Hsp70 [41]. Altogether, our findings and those of others suggest that the function of Hop may vary across species and may also depend on the prevailing cellular physiological conditions.

Findings on the oligomeric status of Hop have remained controversial as independent studies have reported it to be either monomeric [32] or dimeric [33], or largely monomeric but forming also weak dimers [34]. In the current study, we sought to establish the oligomeric status of PfHop. Based on size-exclusion chromatography and multi-angle static light scattering, we conclude that PfHop occurs as a monodisperse monomer (Figure 1). However, a dimeric fraction is observed in non-reducing conditions, indicating unspecific disulphide bridge formation, which would explain some of the previous data [12]. On the other hand, human Hop has been reported to form dimers [42; 33; 43], but it should be confirmed, whether these are of functional significance. In addition, Hop has also been suggested to form elongated monomers, which are difficult to resolve using gel filtration [32]. Indeed, the low-resolution solution structure determined by SAXS shows that PfHop is highly elongated with folded domains organised like beads on a string. This is consistent with the predicted concave nature of its predominantly α-helical TPR motifs [19]. TPR motifs of human Hop have been described to occur as grooves into which the C-terminal EEVD motifs of Hsp90 and Hsp70 bind in extended form [30]. Our SAXS data for PfHop fits with this proposed model.

Altogether, our findings established that PfHop is an elongated, predominantly α-helical, monomeric, protein. Its heat stability is much lower than that reported for PfHsp70-1, suggesting that its function may be compromised at high temperatures associated with clinical malaria progression.

## Acknowledgements

We thank Drs. Arne Raasakka, Erik Hallin, and Juha Kallio for SRCD and SAXS data collection as well as help with data processing. We acknowledge the KIT light source for provision of instruments at the beamline UV-CD12 of the Institute of Biological Interfaces (IBG2), and we would like to thank the Institute for Beam Physics and technology (IBPT) for the operation and storage ring, the Karlsruhe Research Accelerator (KARA). We would like to acknowledge the Biophysics, Structural Biology, and Screening (BiSS) facilities at University of Bergen for access to the static light scattering instrument. This study has been funded by the Academy of Finland, the Norwegian Research Council, the Sigrid Jusélius Foundation, a grant (L1/402/14–1) provided to A.S. by the Deutsche Forchungsgemeinshaft (DFG) under the theme, “German–African Cooperation Projects in Infectiology”, the Department of Science and Technology/National Research Foundation (NRF) of South Africa equipment grant (UID, 75464) and NRF mobility grant (UID, 92598) awarded to A.S.; T.Z. is a recipient of the NRF Innovation Post-Doctoral fellowship UID, 111989 and African–German Network of Excellence in Science junior researcher grant.

